# Revealing unidentified heterogeneity in different epithelial cancers using heterocellular subtype classification

**DOI:** 10.1101/175505

**Authors:** Pawan Poudel, Gift Nyamundanda, Chanthirika Ragulan, Rita T. Lawlor, Kakoli Das, Patrick Tan, Aldo Scarpa, Anguraj Sadanandam

## Abstract

Cancers are currently diagnosed, categorised, and treated based on their tissue of origin. However, how different cellular compartments of tissues (e.g., epithelial, immune and stem cells) are similar across cancer types is unknown. Here we used colorectal cancer subtypes and their signatures representing different colonic crypt cell types as surrogates to classify different epithelial cancers into five heterotypic cellular (heterocellular) subtypes. The stem-like and inflammatory heterocellular subtypes are ubiquitous across epithelial cancers so capture intrinsic, tissue-independent properties. Conversely, well-differentiated/specialized goblet-like/enterocyte heterocellular subtypes differ across cancer types due to their colorectum-specific genes. The transit-amplifying heterocellular subtype shows a dynamic range of cellular differentiation with shared common pathways (Wnt, FGFR) in certain cancer types. Importantly, this approach revealed previously unrecognised heterogeneity in pancreatic, breast, microsatellite-instability enriched and *KRAS* mutation-dependent cancers. Immune cell-type differences are common and useful for patient stratification for immunotherapy. This unique approach identifies cell type-dependent but tissue-independent heterogeneity in different cancers for precision medicine.

## Introduction

Tumors have traditionally been classified into histological subtypes primarily based on their anatomical site of origin (organs or tissues). Although certain mutations (e.g., *TP53*, *PIK3CA*), somatic copy number aberrations (e.g., the *MYC* locus), and gene expression patterns (e.g., basal) are common to several cancer types, cancers are usually described according to their histological differences rather than their molecular and cellular similarities. We propose a different approach in which cancers are regarded according to a cell-of-origin/phenotype model based on tissue architecture and molecular similarities: a heterocellular mix of basal/myoepithelial, epithelial/luminal, immune, connective tissue, and stem cells^1^. Systematic molecular mapping of tissue heterotypia across different cancer types helps to appreciate how the molecular correlates of tissue morphology affect the cancer phenotype and expression of therapeutic targets.

Here we test the hypothesis that common heterocellular signatures across different epithelial cancers define consensus tissue-independent subtypes harbouring common cancer pathways and somatic aberrations. Defining consensus heterocellular subtypes provides opportunities for personalised diagnosis and treatment across the entire epithelial cancer spectrum by establishing common drug targets that may be potentially exploited by repositioning existing drugs. We achieved this by constructing consensus heterocellular subtypes based on our colorectal cancer (CRC) gene expression subtypes (CRCassigner; enterocyte, goblet-like, inflammatory, stem-like, and transit-amplifying (TA)^2^), since these already represent different cells occupying normal and diseased colonic crypts^2^. Also, the FDA has approved the most drugs for CRC^3^, providing broader scope for rapid drug repositioning into other cancers.

Using CRCassigner gene expression signatures to reclassify other epithelial cancer types, we show that these five heterocellular subtypes are prevalent in other cancers. Consequently, we could characterise immune enrichment, microsatellite instability (MSI), and mutational enrichment in tumours from different epithelial cancers using CRC subtypes as surrogates. The molecular features previously thought to be specific for CRC are present in other cancers and can be exploited to select the most appropriate therapy irrespective of cancer type.

## Results and Discussion

### Presence and variability of CRC subtypes across different epithelial tumours

We sought to reclassify individual epithelial cancers as consensus heterocellular subtypes reflecting the overall tissue/cellular architecture (**Figure 1A**). We started with the CRCAssigner subtypes, since they already segregate tumours according to the cells of the colon crypt. Correlating CRCAssigner signature-based PAM (prediction analysis of microarrays^4^) centroids to gene expression profiles of individuals across a range of 10 epithelial cancer types [CRC^5^ (n=262; as a comparison), gastric^6^ (GC/GSE15459; n=182), pancreas^7^ (pancreatic adenocarcinoma and its variants, PC; n=96), lung adenocarcinoma^5^ (LAUD; n=351), endometrial^5^ (UCEC; n=370), breast^5^ (n=835; BRCA), bladder^5^ (BLCA; n=122), head and neck^5^ (HNSC; n=303), ovarian^8^ (OV/GSE9891; n=177), kidney^5^ (KIRC; n=480), and lung squamous carcinoma^5^ (LUSC; n=257)] reclassified each cancer into consensus heterocellular subtypes (**Figure 1B and Supplementary Table 1A-F**) and “mixed/undetermined” tumours that did not classify as a heterocellular subtype. All subtypes were present in all ten cancer types at variable proportions (**Figure 1B**), which we attributed to three factors.

**Figure 1.**
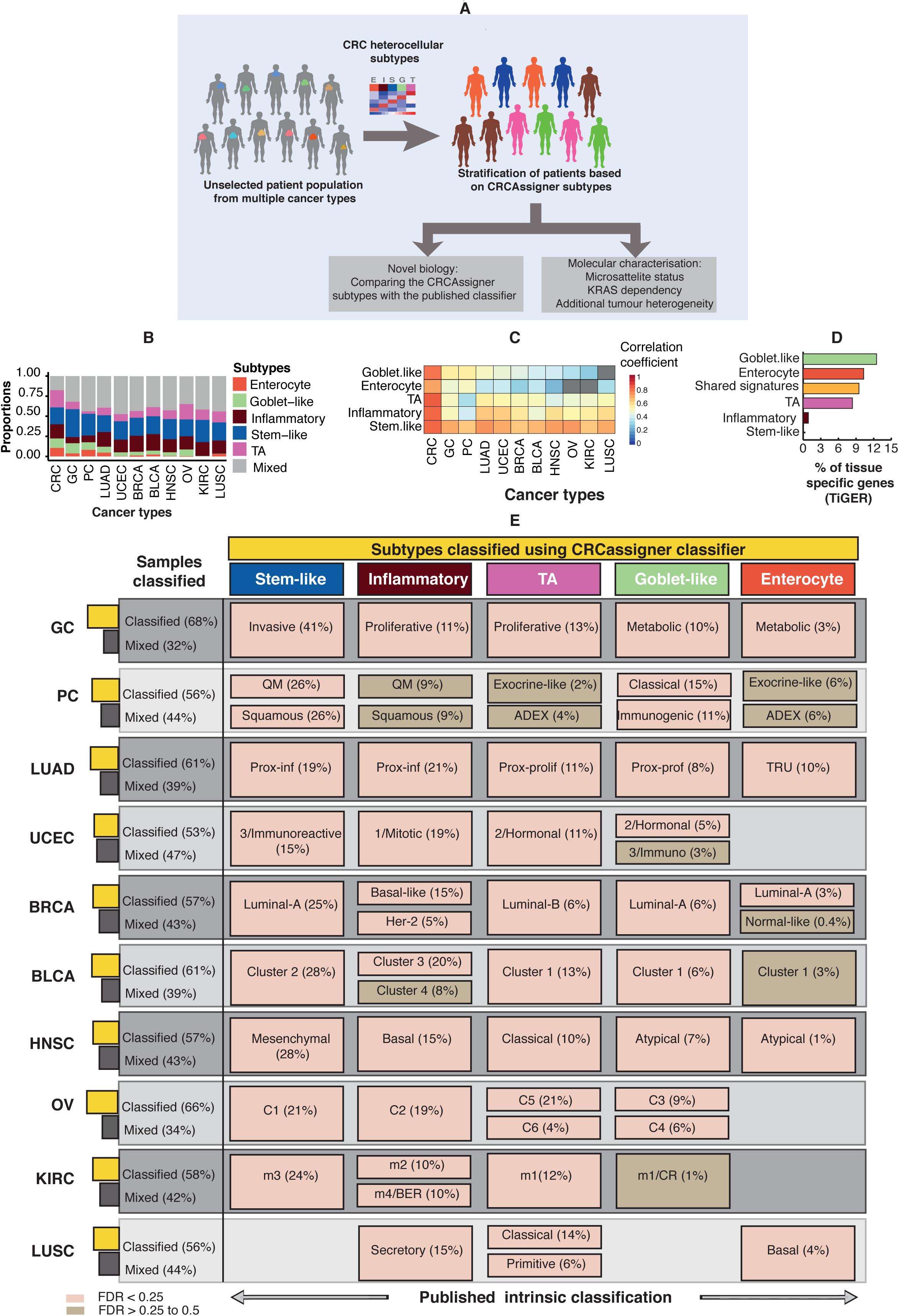
Classification of multiple cancer types into CRC heterocellular subtypes using CRCassigner gene signatures and comparison of CRC heterocellular subtypes with the corresponding intrinsic subtypes. **A.** Schematic of the approach taken to classify multiple cancer types into CRCassigner heterocellular subtypes to understand biological and molecular characteristics. **B.** Proportion of CRC heterocellular subtypes in different cancer types from different data sources [CRC^5^ (Pan-Cancer; n=262), GC^6^ (GSE15459; n=182), PC^7^ (ICGC; n=96), LUAD^5^ (Pan-Cancer; n=351), UCEC^5^ (Pan-Cancer; n=370), BRCA^5^ (Pan-Cancer; n=835), BLCA^5^ (Pan-Cancer; n=122), HNSC^5^ (Pan-Cancer; n=303), OV^8^ (Pan-Cancer; n=177), KIRC^5^ (Pan-Cancer; n=480), LUSC^5^ (Pan-Cancer; n=257). **C.** Heatmap showing correlation coefficients comparing CRCassigner PAM centroids and median values of genes across samples within each subtype and cancer type. Grey colour represents the subtypes (in each cancer type) with less than 2 samples. **D.** Percentage of colon-specific genes (from TiGER^15^) present in the CRCassigner gene signature. Shared signatures represent the genes associated with two or more subtypes. **E**. Comparison of CRC heterocellular subtypes with the intrinsic subtypes of the cancer types present in panel B using hypergeometric test-based FDR values. The cancer types in panels B and E are ordered based on the correlation coefficient of goblet-like subtypes in panel C. Quasi-mesenchymal (QM), aberrantly differentiated endocrine exocrine (ADEX), terminal respiratory unit (TRU), C1 (stroma rich), C2 (immune rich), C3 (secretory type), C4 (high grade), C5 (Wnt high), C6 (deregulated Wnt), immuno-reactive (IR), chromatin remodelling (CR), base excision repair (BER).

First, some variability arose as a consequence of the specific cancer type^9^: taking the mixed/undetermined subtype as an example, <32% gastric cancers (considering four datasets: GSE15459, GSE34942, GSE35809 and TCGA) and 47% UCEC were mixed/undetermined (**Supplementary Figure 1 A-C and Supplementary Table 1A-F**) potentially due to function, location, embryonic origin, intra-tumoural heterogeneity, or normal or other cell type contamination. Different cancers were variably similar to CRCs (**Figure 1C**). For instance, gastric cancers were similar to CRCs for the goblet-like and enterocyte subtype, followed by pancreatic and lung adenocarcinomas (**Figure 1C, Supplementary Figure 1D and Supplementary Table 1G**), mirroring the presence of mucin-secreting cells in colorectal, gastric, pancreatic, and lung adenocarcinomas. In contrast, the inflammatory and stem-like subtypes were similar across different cancer types (**Figure 1C and Supplementary Figure 1D**), suggesting that the immune and stem cell compartments are similar in different cancers.

Second, some variability may be dataset specific, so we validated the presence of the CRC heterocellular subtypes in other gastric cancer^6,10^ (three datasets; n=363), pancreatic cancer^11^ (n=36), colorectal^12^ (n*=288*), ovarian^5^ (n*=259*), lung^13^ (LAUD and LUSC; n*=168*), and breast^14^ (n=104) datasets (**Supplementary Figure 1F and Supplementary Table A-F**). The enterocyte subtype was not present in ovarian^5^ or kidney^5^ cancers and was present only in a small proportion of other cancers, probably because enterocytes are highly specialised, terminally differentiated, and present only in the small intestine and stomach (**Supplementary Figure 1F and Supplementary Table 1E-F**). For the same reason, the goblet-like subtype was uncommon in kidney (0.6%) and lung squamous cell cancers (0.4%; **Supplementary Table E-F**). However, at least four CRC heterocellular subtypes were present in the majority of cancer types.

Third, tissue-specific genes might contribute to heterocellular subtype variability across cancer types, so we shrunk the CRCassigner signature into 564 (CRCassigner-564) genes specific for each subtype using PAM centroid scores (**Supplementary Table 1H**) and compared them to colon-specific genes (n=198; present in expressed sequence tag; **Supplementary Table 1I**) from the tissue-specific database^15^ (TiGER). The goblet-like subtype had the highest percentage of colon-specific genes (12.5%, n=6) followed by enterocyte (10.3%, n=8) and TA (8.4%, n=13) subtypes (**Figure 1E and Supplementary Figure 1E and Supplementary Table 1J**), accounting for the high variability of these subtypes across cancer types (**Supplementary Figure 1F-I**). Conversely, none of the stem-like subtype marker genes and only one from the inflammatory subtype were colon specific. The remaining gene signature (n*=*222) shared by two or more subtypes contained 9% of the colon-specific genes, further contributing to variability. Overall, at least four heterocellular subtypes were similar across cancer types while their proportions varied depending on the functional and tissue-specific characteristics.

### Comparison of heterocellular subtypes to known intrinsic subtypes in multiple cancer types

Tissue-specific cancers are now commonly subdivided into different heterocellular subtypes. To further understand which heterocellular subtypes correspond to which intrinsic cancer subtypes, we compared heterocellular subtypes with the original published/known subtypes for different cancers. There were significant associations for ten cancer types (chi-squared test; p<0.0001, **Figure 1E and Supplementary Table 1K**). **Figure 1E** summarises the systematic associations between intrinsic subtypes with heterocellular subtypes and the characteristics of CRC subtypes. Overall, in the majority of cases, the intrinsic subtypes with stemness, mesenchymal, and stroma-rich characteristics and poor patient prognosis corresponded to the stem-like heterocellular subtype and those with immune enrichment and intermediate prognosis corresponded to the inflammatory subtype (**Supplementary Table 1L-N**). In contrast, those with differentiated and secretory phenotypes corresponded to the goblet-like subtype.

Interestingly, the TA heterocellular subtype across cancers was similar to intrinsic subtypes with a dynamic differentiation potential and unique pathway enrichment and characteristics. For instance, Wnt signature-high C5 and C6 ovarian cancer subtypes were significantly associated with the TA subtype with high Wnt signalling. The exocrine-like intrinsic pancreatic cancer subtype and classical HNSC subtype with TA subtype gene profiles showed xenobiotic metabolic pathway enrichment^16^. In bladder cancer, the “cluster I” subtype^17^ with papillary histology and *FGFR3* aberrations was significantly enriched for the *FGFR3*-high TA subtype. The “m1” kidney cancer subtype was enriched for the TA subtype representing a potential chromatin-remodelling process (**Figure 1E**).

Therefore, although some associations between heterocellular and intrinsic subtypes were variable across cancer types, the core tissue heterotypia described by heterocellular subtypes was reflected in the existing molecular subtypes, i.e., epithelial/mesenchymal, differentiated/undifferentiated/stemness, or luminal/basal/stemness. Given that heterocellular subtypes did not perfectly match existing subtypes, we further analysed what emergent biology was discernible in different cancer types given the heterocellular classification.

### Pancreatic cancer subtypes

We previously described three intrinsic pancreatic adenocarcinoma gene expression (PDAssigner^18^) subtypes: classical, quasi-mesenchymal (QM), and exocrine-like that overlapped with Bailey et al.’s recently published subtypes^7^: the pancreatic-progenitor subtype, with classical subtype overlap; squamous, with QM-PC overlap; aberrantly differentiated endocrine exocrine (ADEX), with exocrine-like overlap; and an immunogenic subtype, a subset of classical subtype^18^ (**Supplementary Figure 2A-B**). Using the PDAssigner training data set (GSE15471^11^, **Supplementary Figure 2C**) and ICGC (Bailey’s, **Figure 2A**) datasets, the differentiated classical PC subtype was associated with the goblet-like subtype **(Figure 2B and Supplementary Figure 2C**). Furthermore, all PC precursor samples with intraductal papillary mucinous neoplasms with invasion (a histological subtype) represented 33% of the goblet-like PC subtype (**Supplementary Figure 2D-F and Supplementary Table 2A-B**) suggesting that one pathway of PC development occurs through goblet-like subtype precursor. The pancreatic-progenitor subtype was more heterogeneous and not exclusively associated with a specific heterocellular subtype (**Figure 2B and Supplementary Figure 2G**), similar to luminal-A breast cancer described below.

**Figure 2.**
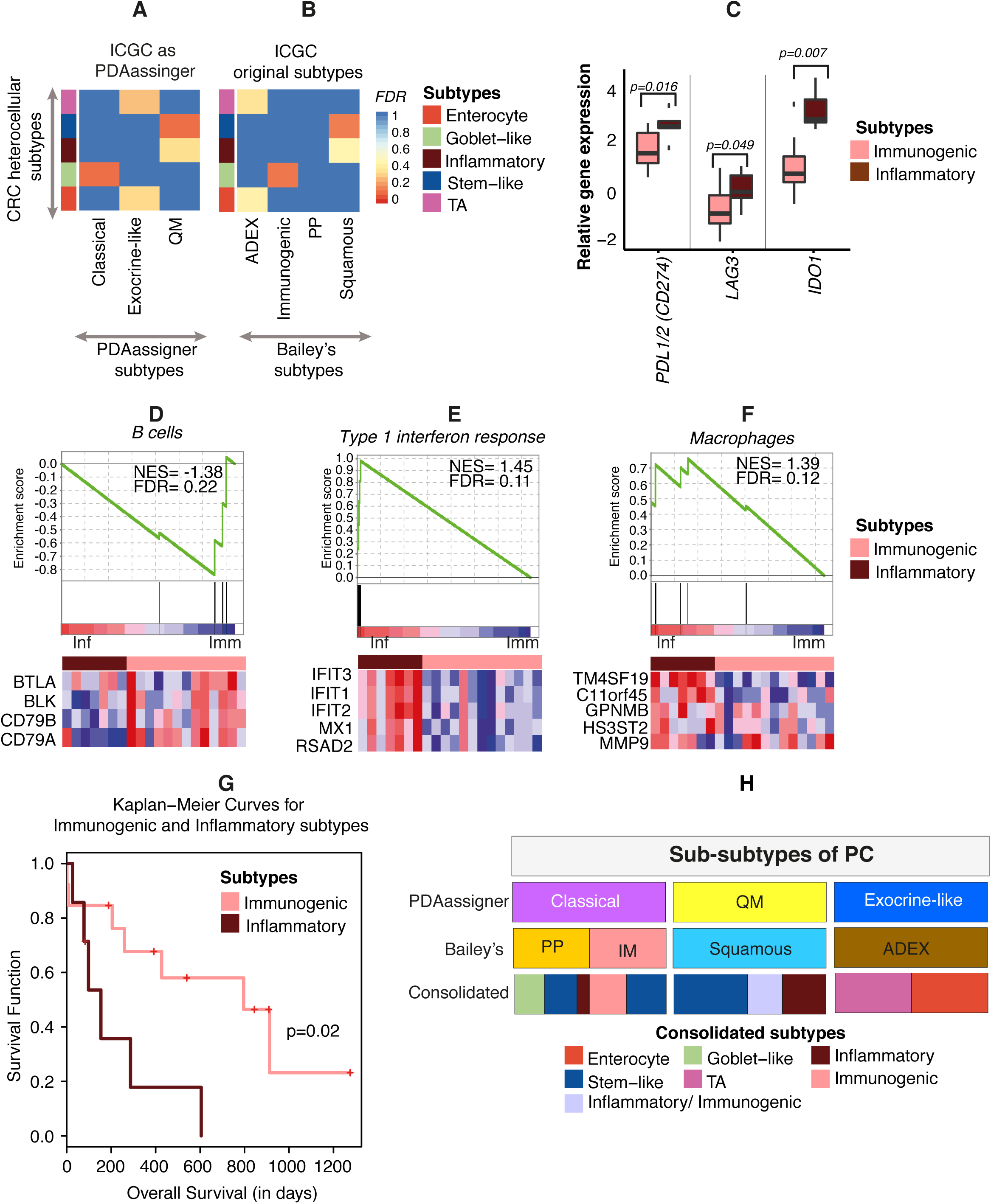
Comparison of intrinsic pancreatic subtypes with CRC heterocellular subtypes and their molecular phenotypes. **A-B.** Heatmap showing hypergeometric test-based FDR values comparing CRC heterocellular subtypes (y-axis) with (A) PDAassigner^18^ subtypes (x-axis) and (B) Bailey’s subtypes^7^ (x-axis) using ICGC^7^ (n=54; removing the mixed/undetermined samples) data in both cases. **C.** Box plot showing differences in the expression of checkpoint immune genes *IDO1*, *PDL1/2 (CD274)*, and *LAG3* between immunogenic (n=13) and inflammatory (n=7) PC subtypes. **D-F.** GSEA analysis (using published^23^ immune markers) showing enrichment of B cells, type 1 interferon response, and macrophages between inflammatory and immunogenic PC subtypes. **G.** Kaplan-Meier survival curve showing significant prognostic (overall survival) difference between immunogenic (n=13) and inflammatory (n=7) PC subtypes. **H**. Summary of the PC subtypes from multiple studies as consolidated subtypes.

In contrast, good prognosis exocrine-like/ADEX PC subtypes were associated (**Figures 2B, Supplementary Figure 2C and Supplementary Table 2A-B**) with good prognosis TA and intermediate prognosis enterocyte CRC subtypes, which were similar enough to form a consensus subtype^19^. Since normal TA cells originate from colon crypt stem cells and are a heterogeneous mixture of well and poorly differentiated cells, this raises the question of whether the exocrine-like subtype originates from acinar cells with transit-amplifying and/or intermediate characteristics (**Supplementary Figure 2D and F**).

The poor prognosis and highly glycolytic QM-PC subtype^18,20^ was associated with poor prognosis stem-like or intermediate prognosis inflammatory heterocellular subtypes (**Figures 2A and Supplementary Figure 2C**). Nevertheless, increased mesenchymal and stromal characteristics are likely to be common to stem-like and inflammatory heterocellular subtypes (**Figure 2C and Supplementary Figure 2H**). In addition, CRC inflammatory-specific genes, which represent mainly innate immune characteristics, were enriched in 30% of pancreatic cancers, validating the presence of the inflammatory subtype in pancreatic cancer. However, a subset of classical/pancreatic-progenitor subtype PCs also showed inflammatory characteristics (**Supplementary Figure 2I and Supplementary Table 2C**) suggesting that inflammation can be independent of intrinsic subtype; the cause of this requires further study.

In contrast, Bailey’s immunogenic PC subtype (analysed from the ICGC dataset) was not associated with the inflammatory subtype, rather primarily with the goblet-like subtype (**Figure 2B**). Hence, we further characterised differences in immune cells and biomarkers between immunogenic and inflammatory subtypes. There were significant increases in *IDO1*, *LAG3*, and *PDL1/2(CD274*) expression in the inflammatory (n=7) subtype compared to the immunogenic subtype (n=13) (**Figure 2C and Supplementary Table 2D**), suggesting that inflammatory subtype patients may respond well to immune checkpoint therapies compared to immunogenic subtype patients. There was also significant enrichment of naïve B cell-specific genes in the immunogenic subtype compared to significantly enriched type-1 interferon and macrophage-specific genes in the inflammatory subtype (**Figure 2D-F and Supplementary Table 2E-F**). Moreover, inflammatory subtype pancreatic cancer patients had shorter overall survival than immunogenic patients (**Figure 2G**). However, the clinical outcomes of that subset of pancreatic cancer samples containing both inflammatory and immunogenic signatures were unclear due to insufficient data (**Supplementary Figure 2I**).

Overall, our approach unrevealed heterogeneity in pancreatic cancers and at least six different subtypes: goblet-like (part of classical/pancreatic-progenitor); stem-like (part of QM-PDA/squamous and classical); immunogenic (mainly part of classical subtype); inflammatory (mainly from QM-PDA/squamous and a small proportion from pancreatic-progenitor); and TA and enterocyte (exocrine-like/ADEX; **Figure 2H and Supplementary Figure 2I**).

### Gastric cancer subtypes

We next compared our heterocellular subtypes to the intrinsic gastric cancer subtypes defined by Lei et al.^*6*^ (**Figure 3A, Supplementary Figure 3A-B and Supplementary Table 1L-N**). The invasive and stem-like subtypes were significantly associated; both had increased cancer stem cell-like properties with mesenchymal characteristics^2,6^. The gastric metabolic subtype was associated with the enterocyte and goblet-like differentiated gastric cancer subtypes and was enriched for metabolic/digestion-associated genes and differentiated, “normal” gastric mucosa-like features^6^. The proliferative subtype was heterogeneous and significantly associated with the inflammatory and TA heterocellular subtypes. Additionally, these heterocellular subtypes were also prognostic, such that the stem-like, TA, and inflammatory subtypes had better survival with surgery alone compared to adjuvant (surgery+) 5-FU treated patients (**Figure 3B and Supplementary Figure 3D**). To better understand this molecular heterogeneity, we compared our heterocellular subtypes to four TCGA integrative stomach adenocarcinoma or gastric cancer (STAD) subtypes^10^ (**Supplementary Figure 3C**). As expected, the inflammatory heterocellular subtype was significantly associated with the microsatellite-enriched (MSI) and Epstein-Barr virus (EBV) STAD subtypes known to have prominent immune cell infiltrates, while the TA heterocellular subtype corresponded to the chromosomal instability (CIN) STAD subtype. Similar to the cetuximab-sensitive (CS)-TA sub-subtype with increased EGFR ligands and associated signalling, the CIN STAD subtype showed EGFR amplification and phosphorylation (PY1068), suggesting the potential for anti-EGFR targeting. Finally, the genomically stable TCGA STAD subtype was associated with the stem-like and enterocyte heterocellular subtypes. Overall, the CRC heterocellular subtypes and intrinsic gastric cancer subtypes are associated and possess similar biological characteristics.

**Figure 3.**
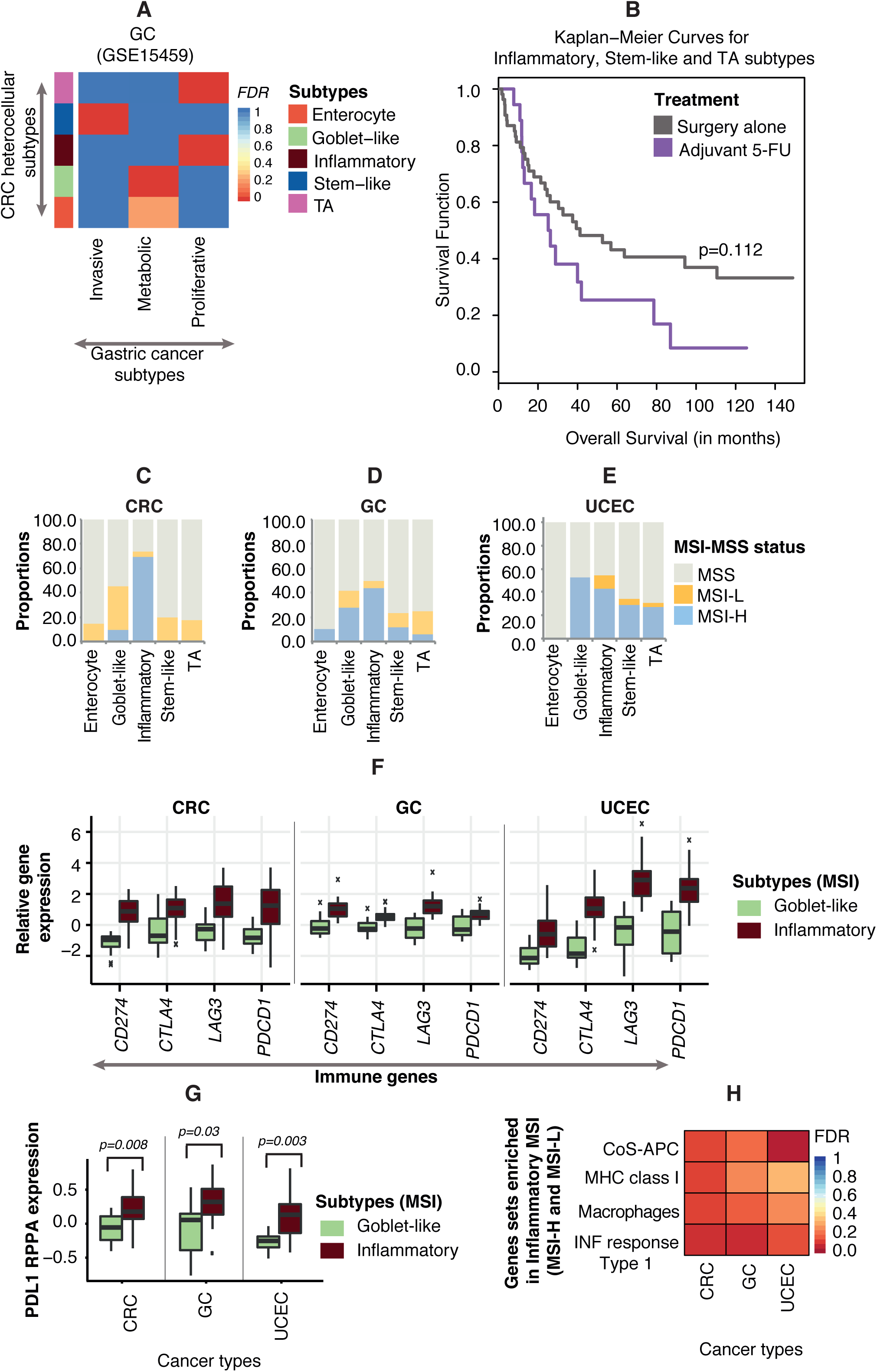
Characterising MSI and MSS phenotypes in different cancer types using heterocellular subtypes. **A.** Heatmap showing hypergeometric test-based FDR values comparing CRC heterocellular subtypes (y-axis) with intrinsic gene expression subtypes from Lei et al.^*6*^ in gastric cancer datasets (GSE15459; n=124; after removing mixed/undetermined samples). **B**. Kaplan-Meier survival curve showing the overall survival difference between surgery (n=57) and adjuvant 5-FU (n=18) treatment groups in combined data (GSE15459 and GSE34942) consisting of stem-like, inflammatory, and TA subtypes. **C-E.** Bar graphs showing the proportion of MSI-L, MSI-H, and MSS samples in different CRC heterocellular subtypes in (C) colorectum^28^ (CRC), (D) gastric^10^ (GC) and (E) endometrium^29^ (UCEC). The MSI status of the three cancer types is taken from the respective TCGA papers^28,10,29^. **F.** Box plot showing significant (p<0.001) differences in the expression of checkpoint immune genes *PDL1/2 (CD274*) *CTLA4*, *LAG3,* and *PDCD1* between MSI samples in goblet-like and inflammatory subtypes in CRC, GC, and UCEC. **G.** Box plots showing differences in the expression of checkpoint immune marker *PDL1* between MSI-enriched goblet-like and inflammatory subtypes from CRC, GC, and UCEC, as assessed using RPPA^30^ data. **H.** Heatmap showing the enrichment of immune cells (co-stimulation APC, MHC class 1, macrophages, type 1 interferon response) between MSI-enriched goblet-like and inflammatory subtypes from CRC, GC, and UCEC, as assessed using GSEA^31^ and published^23^ immune markers.

### Heterogeneity in MSI cancers dictated by heterotypic subtypes and the immune microenvironment

We sought to explore whether our heterocellular subtypes show similar therapeutic and molecular features to their parent subtypes. We previously showed that inflammatory subtype CRCs were significantly enriched for microsatellite instable (MSI)^2^, so we reasoned that similar enrichment for MSI would be present in other cancer types (**Figure 3C-E and Supplementary Table 3A-I**). Similar to CRC (TCGA) inflammatory samples with greater than 75% MSI (both MSI-H and MSI-L, **Figure 3C**), the inflammatory subtype in gastric (stomach; TCGA STAD, **Figure 3D**) and endometrial (TCGA UCEC, **Figure 3E**) cancers had >40% MSI. The goblet-like subtype had about 40% MSI (mostly MSI-L) and, similarly, the goblet-like cancers from both TCGA STAD and UCEC showed the second highest proportion of MSI samples.

Multiple studies^21,22^ have suggested that MSI tumours could be susceptible to immune checkpoint blockade, so we compared the expression of immune genes between the MSI samples from goblet-like and inflammatory subtypes across the three cancer types. Interestingly, inflammatory MSI tumours showed significantly (p<0.001) higher expression of *PDL1/2(CD274)*, *CTLA4*, *LAG3*, and *PDCD1* genes compared to goblet-like MSI tumours in all three cancer types (**Figure 3F**). Additionally, inflammatory MSI tumours from all three cancer types showed increased protein expression of *PDL1* compared to that of the goblet-like subtypes, probably indicating that inflammatory MSI patients may derive greater benefit from anti-PDL1 therapy (**Figure 3G**). To further understand the immune cell type and pathway differences between goblet-like and inflammatory MSI tumours, we used published immune gene markers^23^ and performed GSEA analysis (**Figure 3H and Supplementary Figure 3M**). Inflammatory MSI tumours showed enrichment of genes associated with co-stimulation of APC (CoS-APC), MHC class I, type I interferon (INF) responses, and macrophages (**Supplementary Table 3J-L**) compared to goblet-like MSI tumours. Overall, our observations indicate that MSI tumours are heterogeneous, so PDL1 or other immune checkpoint therapies may fail in practice if this heterogeneity is not appreciated. With careful selection of patients based on inflammatory vs. goblet-like differences in MSI tumours, however, immune checkpoint therapy responders may be identified.

### *KRAS* mutant cancers are heterogeneous and the goblet-like subtype is *KRAS*-dependent

*KRAS* mutations are characteristic of CRC and negatively predict anti-EGFR responses. We previously showed that CMS3 (equivalent to the goblet-like CRCassigner subtype) was enriched for *KRAS* mutations^19^. Interestingly, the goblet-like subtype was enriched (76%) for *KRAS* mutations in the TCGA CRC dataset (**Figure 4A and Supplementary Table 4A-B**) compared to <40% for other subtypes. Similarly, lung adenocarcinomas harbour *KRAS* mutations and, accordingly, 50% of the goblet-like TCGA LUADs were *KRAS* mutated (**Figure 4B and Supplementary Table 4C-D**). *KRAS* mutations are present in 89% of pancreatic cancers so mutation status alone is non-discriminatory (**Figure 4C and Supplementary Table 4E-F**), so we assessed if the *KRAS* dependency reported by Singh et al.^24^ and ourselves^18^ was similar in all three cancer types using the *KRAS* dependency signature^24^ and nearest template prediction method^25^. The *KRAS* dependency signature was enriched in all three cancer types (FDR<0.02 for CRC, FDR<0.18 for LUAD, and FDR<0.36 for PC; **Figure 4D-F and Supplementary Table 4G-L**) in the goblet-like subtype. The enterocyte subtype was also enriched for *KRAS* dependency in TCGA CRC and PC samples, while the stem-like and inflammatory subtypes were *KRAS* independent. To examine whether *KRAS* dependent/independent tumours share a common mechanism, we selected *KRAS*-dependent goblet-like and *KRAS*-independent stem-like samples and performed pathway analysis using GSEA across the three cancer types. The epithelial-mesenchymal transition (EMT) pathway was significantly (FDR<0.1) enriched in *KRAS*-independent groups (**Figure 4G and Supplementary Table 4M-O**), consistent with previously reported findings *in vitro*^24^ that *KRAS*-independent cells have more EMT-like phenotypes. Therefore, any therapy that is effective in goblet-like-dependent or stem-like-independent *KRAS* mutation samples may be applicable to the other two cancers with the same profiles.

**Figure 4.**
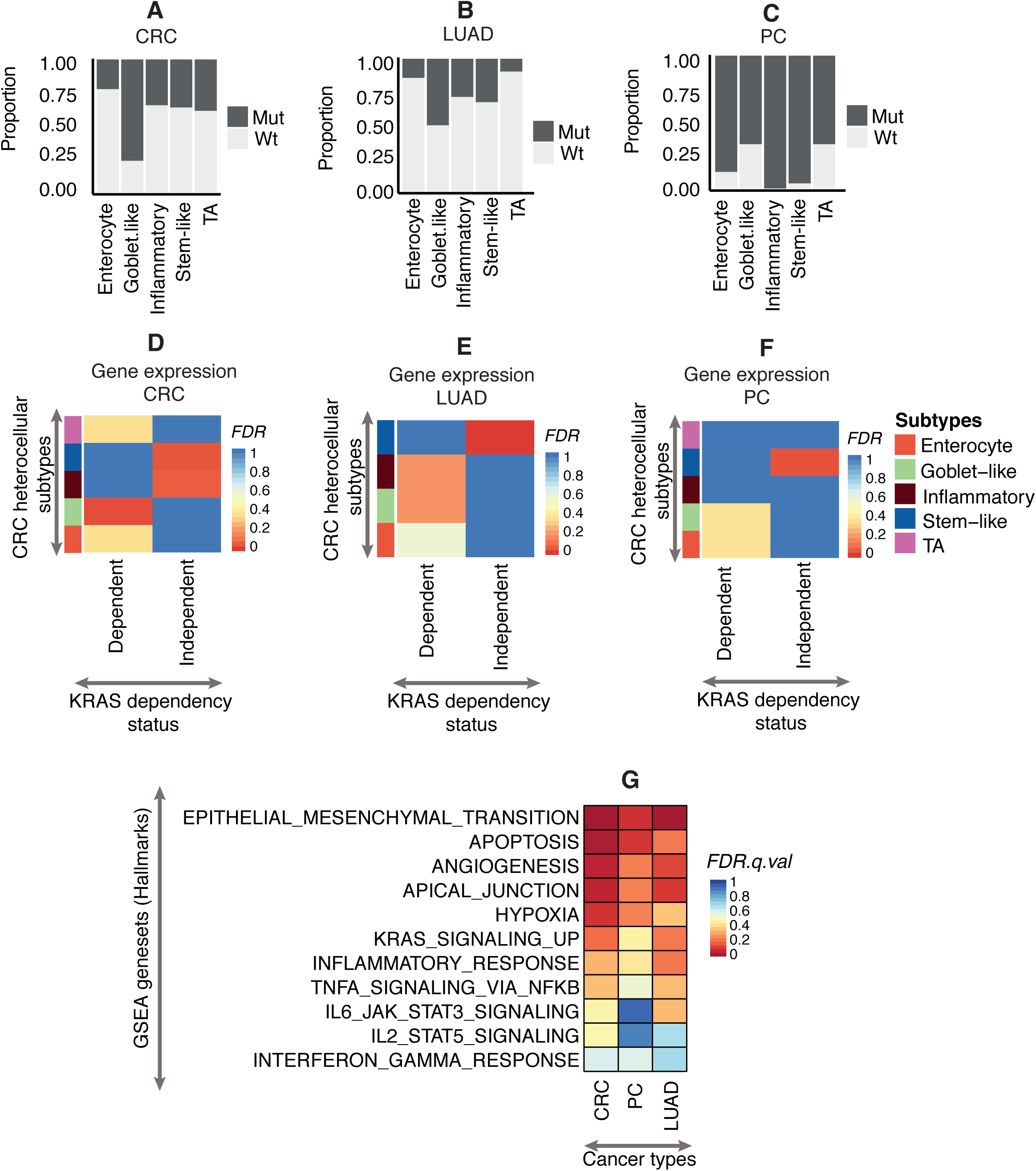
Characterising *KRAS* dependency status in *KRAS*-mutant samples from three different cancers. **A-C.** Proportions of *KRAS*-mutant and wild type samples in CRC heterocellular subtypes from (A) CRC^28^ (61 mutant and 90 wild type), (B) LUAD^32^ (37 mutant and 94 wild-type), and (C) PC^7^ (48 mutant and 6 wild-type). **D-F.** Heatmap showing hypergeometric test-based FDR values comparing *KRAS*-mutant samples (in CRC heterocellular subtypes) with the *KRAS* dependency^24^ status of the corresponding samples predicted using the NTP^25^ method in (D) CRC, (E) LUAD, and (F) PC. **G.** Enrichment of hallmarks^33^ gene-sets in *KRAS*-independent stem-like subtypes compared to the *KRAS*-dependent goblet-like in CRC, PC and LUAD.

### Luminal breast cancer heterogeneity associated with progenitor cells

The basal-like breast cancer and HER2-enriched subtypes^26^ were significantly associated with the inflammatory subtype (**Figure 5A, Supplementary Figure 5A and Supplementary Table 1L-N**), with basal-like cases similar to equivalent subtypes in HNSC and bladder cancer (**Supplementary Figure 3F and 3H**). The luminal B subtype was significantly associated with the TA subtype, suggesting that luminal B cancers might have a transitional phenotype between stem and differentiated cell types, like in the colonic crypt.

**Figure 5.**
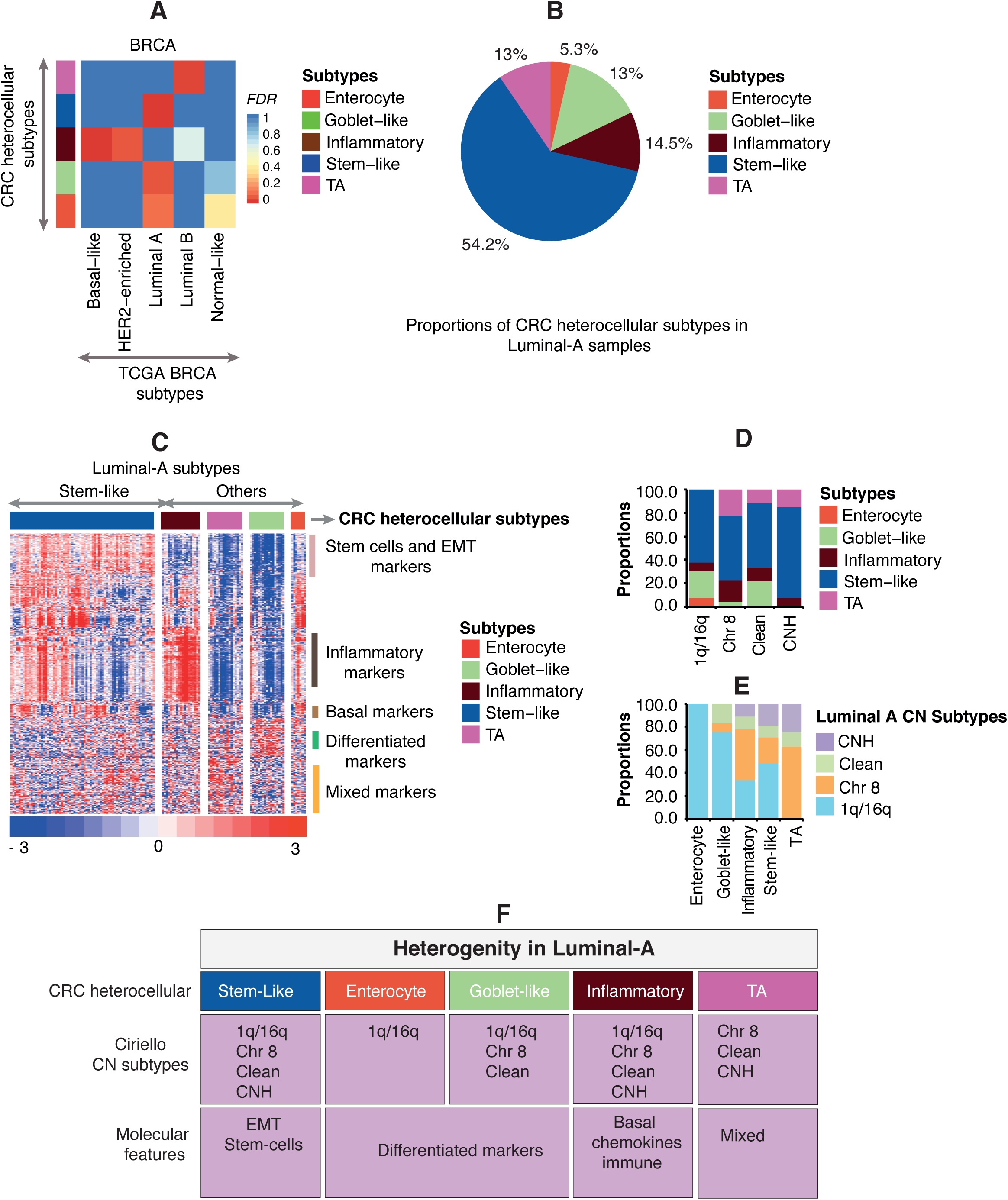
Characterisation of heterogeneity in luminal A breast cancers. **A.** Heatmap showing hypergeometric test-based FDR values comparing CRC heterocellular subtypes (y-axis) with intrinsic gene expression BRCA subtypes^26^ (x-axis). **B.** Pie chart showing proportions of different CRC heterocellular subtypes in luminal A BRCA samples. **C.** Heatmap showing the expression of the top highly variable (SD>1.5) selected marker genes between stem-like (n=71) and other (n=60) subtypes within the luminal A BRCA subtype. **D-E.** Bar plot showing the proportion of (D) CRC heterocellular subtypes in Ciriello et al.^27^ subgroups of luminal A and (E) vice versa. F. Summary of the heterogeneity of luminal A subtypes identified using the CRCassigner signature.

However, of most significance was heterogeneity in the well-characterised luminal A subtype, which was not only associated with the differentiated goblet-like and enterocyte subtypes but also with the poorly differentiated stem-like heterocellular subtype in both BRCA (TCGA and GSE42568) datasets (**Figure 5A; Supplementary Figure 5A**). Luminal A samples were represented by all the heterocellular subtypes in the TCGA BRCA dataset: 54.2% stem-like, 14.5% inflammatory, 13% goblet-like, 13% TA, and 5.3% enterocyte (**Figure 5B**). GSEA revealed enrichment of stem cell genes in stem-like luminal A samples (**Figure 5C; Supplementary Figure 5C and Supplementary Table 5A**), suggesting that stem-like luminal A cancers may have a luminal progenitor cell of origin. Survival analysis showed that stem-like luminal A patients have a trend to poor recurrence-free survival compared to the other luminal-A subtypes (**Supplementary Figure 5B**), however the data is not significant.

Finally, we compared our heterocellular luminal A subtype classification with Ciriello et al.’s DNA copy number and mutation luminal A subtypes^27^ (**Figure 5D-E and Supplementary Table 5B-D**). The well-differentiated enterocyte and goblet-like subtype samples were primarily the 1q/16q Ciriello subtype and TA subtype samples were primarily Ciriello Chr 8 cancers. As expected, the stem-like and inflammatory subtype samples were heterogeneous and represented all four Ciriello subtypes, and the stem-like subtype had a scrambled genome such that 77% belonged to Ciriello copy number high subtype. Overall, these results confirm the heterogeneity in luminal A cancers and provide potential pathogenetic mechanisms from different tissue compartments (**Figure 5F**).

## Conclusion

Here we successfully classified multiple cancer types into five tissue-independent CRC subtypes based on the main cell types present in the colonic crypt. While the majority of cancer types were significantly classified into stem-like and inflammatory subtypes, not all cancer types were readily classified into enterocyte and goblet-like subtypes due to their colon specificity. It is unknown why a certain subset of tissue-independent genes are not dynamically expressed in cancers that are physically or functionally unrelated to the colorectum, but it might be because the majority of CRCassigner genes were those normally associated with normal gastrointestinal-related organs.

This reclassification approach serves to characterise additional heterogeneity in different cancer types, for example three new subtypes in PC. Similarly, we were able to identify additional gene expression-based heterogeneity in the luminal A breast cancer subtype and heterogeneity in MSI or *KRAS* mutant cancers using CRC subtypes as surrogates. MSI in CRC, GC, and UCEC has previously unidentified heterogeneity not only defined by inflammatory and goblet-like genes but also stemness. Finally, *KRAS* mutations in CRC, PC, and lung cancer are heterogeneous and have different dependencies in different subtypes. Overall, these sub-subtypes and the discovery of further heterogeneity defined by consensus heterocellular signatures are useful for diagnostic stratification and to personalise new and existing treatments to shared molecular features in different cancer types.

